# Algorithms for the discovery of cis-eQTL signals in woody species: the vine (*Vitis vinifera* L.) as a study model

**DOI:** 10.1101/2021.07.06.450811

**Authors:** Pedro José Martínez-García, J Mas-Gómez, Jill Wegrzyn, Juan A. Botía

**Affiliations:** Department of Plant Breeding, Centro de Edafología y Biología Aplicada del Segura, Consejo Superior de Investigaciones Científica (CEBAS-CSIC), P.O. Box 164, 30100 Espinardo, Spain; Department of Ecology and Evolutionary Biology, University of Connecticut, CT, 06269, Storrs, USA; Department of Neurodegenerative disease, University College London, London WC1N 3BG, UK; Departamento de Ingeniería de la Información y las Comunicaciones, Universidad de Murcia, Murcia 30100, Spain

**Keywords:** grapes, gene expression, genetic variant, SNP effect

## Abstract

Expression quantitative trait loci (eQTLs), are associations between genetic variants, such as Single Nucleotide Polymorphisms (SNPs), and gene expression. eQTLs are an important tool to understand the genetics of gene expression of complex phenotypes. eQTLs analysis are common in human studies and in model species such as mice, rats and yeast but are very scarce in wood crop species such as fruit trees or grapevines. In this study a comprehensive bioinformatic pipeline has been carried out using genomics and expression data from 10 genotypes of grape, which has been used as model species. As a result of this study a total of 10,618 genetic variants that regulate gene expression levels of 525 genes were detected. A 78% of them, 411, received a functional annotation from UniProtKB or DAVID, between the annotated protein-coding genes are Germin-like proteins (GLPs), auxin-regulatory factors, GRFS, ANK_REP_REGION domain-containing protein, Kinesin motor domain-containing protein or RPP13like protein 2(LOC100852873). This new inventory of cis eQTLs influencing gene expression during the ripening of fruits of *Vitis vinifera* L. will be an important resource for future research to understand the mechanistic basis for variation in gene regulation in this species. In the future, this methodology may be applied to other woody species, once the necessary databases are generated.

## Introduction

A phenotype or trait with suspected or known genetic behaviour that does not follow to classical Mendelian heritage is generally called as a complex genetic trait [1]. Many, if not most, of important characters to animal, plant and human research, such as morphological, life history, behavioral traits, as well as many human diseases such as cancer and diabetes are genetically complex. The concept *complex* is reserved for phenotypes that are controlled by many genes (polygenic) as well as environmental conditions. Complex traits also encompass a type of phenotypes that being discontinuous (or discrete) are only expressed when the effects of many genes and/or environmental conditions reach minimum threshold. Susceptibility to any important infectious diseases to mammals health, such as SARS-CoV-2, falls in this class as well. To know the procedures that generate differences in complex traits among individuals, populations, and species has been an essential challenge to evolutionary biology since Darwin and Galton [2]. In addition, the inheritable contribution typically accounts for a considerable proportion of the variation amongst individuals [2]. Understanding of phenotypic differences among individuals hence demands the knowledge of the causes of genetic variation in complex traits.

The systematic dissection of the genetic basis of variation in complex traits has just become achievable over the last ten years [3]. Since 2007, genome-wide association studies (GWAS) have detected association between common genetic variation at thousands of loci and important human diseases [4]. In 2021, the catalog of published GWAS (www.genome.gov/gwastudies) includes 5,106 publications and 161,014 SNPs as of June 8, 2021. Surprisingly, the number of specific mutations identified and demonstrated to be causative has been very low, in comparison with the high numbers of genomic loci underlying complex traits mapped. The two main reasons for the difficulty of understanding GWAS associations are, first that nearby genetic variants are probably inherited together due to co-segregation during meiotic recombination, a phenomenon called as linkage disequilibrium (LD), hindering the identification of the causal variants propping the association. Secondly, it is unknown which cell types are causal to the disease, as the pathophysiology of complex diseases often involves interactions of many cell types [5]. More interestingly, nearly 90% of these trait/disease associated SNPs (TASs) were located outside of protein-coding regions, suggesting a possible role in gene regulation of these associated variants [6]. All this reflects that, in spite of this amount of GWAS, the amount of reports that have explored the underlying mechanisms of the detected associations, meaning the functional effects of GWAS-implicated variants, is orders of magnitude fewer [7, 8].

One of the the biggest challenge in the actual post-GWAS era lies in connecting additional molecular data with these GWAS findings to functionally illustrate the relationships. In this sense, improvements in omics technologies have allowed to explore the impact, on intermediate molecular levels, of the risk variants, like protein abundance, methylation, metabolite levels or gene expression. The combination of current bioinformatic, statistical, and empirical bench-based methods is allowing the downstream elucidation of GWAS-identified trait loci. In this sense, genetical genomics is a very powerful tool to identify expression quantitative trait loci (eQTLs), meaning the genomic loci that control gene-expression differences. Its aim is to improve our understanding on the genetic architecture of disease susceptibility and complex traits. And thus to improve our knowledge of how gene networks are organised within an organism [9]. Almost 50% of trait-associated SNPs have a cis-acting effect on gene expression [10, 11]. According to this, cis-eQTL associations have been performed to effectively detect the novel but weak associations that no reach the genome-wide significance level in GWAS, without the requirement of increasing the size of the sample [12, 13]. Other significant benefit of the detection of eQTLs is that they can provide knowledge about the underlying pathways and the trait mechanism [14]. Different studies have confirm the increasingly importance of resolving the molecular pathways leading to disease, due to the fact that trans-eQTLs can impact downstream disease genes not previously discovered by GWAS [15]. However, a efficient recognition of trans-eQTLs is hard, because trans-eQTL are indirect association with a effect much smaller than a cis-eQTL. As a result of two different theories, the theory of phenotypic buffering and the robustness of biological systems, molecular elements placed further downstream in a pathway have a smaller effect than the elements upstream [16]. To characterize functional genetic variation in humans, a large consortium [17] has allowed the development of the Genotype-Tissue Expression (GTEx) project supported by the NIH Common Fund. This project represents a great resource, having associated a great tissue bank. The presence of this resource, allows researchers to study the liaison between gene expression and genetic variation and other molecular data in a large number of tissues. Studying large amount of tissues or cell types is expensive and laborious being limited to humans and unaffordable for plant species to date such as woody crops. However, eQTL studies are earning popularity in plant genetics mainly due to the fact that they represent a efficient approach to short out the tedious procedure of positional cloning, especially for genes underlying quantitative characters [18]. At the same time, eQTL studies of tree species is a real challenge because of sampling methods [19] since they do not grow easily under controlled environments due to large space requirements. Indeed, few eQTL studies have been conducted in woody species [20, 21, 22, 23, 24, 25, 26]. Some of these studies have been carried out in grapes (*Vitis vinifera* L), the target organism of this study. This crop was one of the first domesticated fruit crops and being now one of the most profitable horticultural crop, with around 8 million ha of vineyard in the world, the majority destined to produce wine. The development of the common grape berry, follows a double sigmoid pattern of growth (Figure 1), with the inception of ripening (véraison) at the beginning of the second development phase, maturation phase [27]. This phase is characterized by some of the most noticeable changes in the grapes: pronounced berry growth, sugar accumulation, decrease of acidity/raise of pH and accumulation/changes in phenolic compounds and aromatic [28, 29]. From a physiological point of view, when seeds are able to germinate grape maturity is reached, which is immediately after véraison. In the published results by [30], the authors found that in general a high increase of DEGs was characterized in the majority of varieties between pre-véraison (PV or Touch) stage and end of véraison (EV or Soft) stage, with a notable decline in the number of differentially expressed genes (DEGs) from end of véraison (EV or Soft) to harvest (H) (Figure 1).

**Figure 1.**
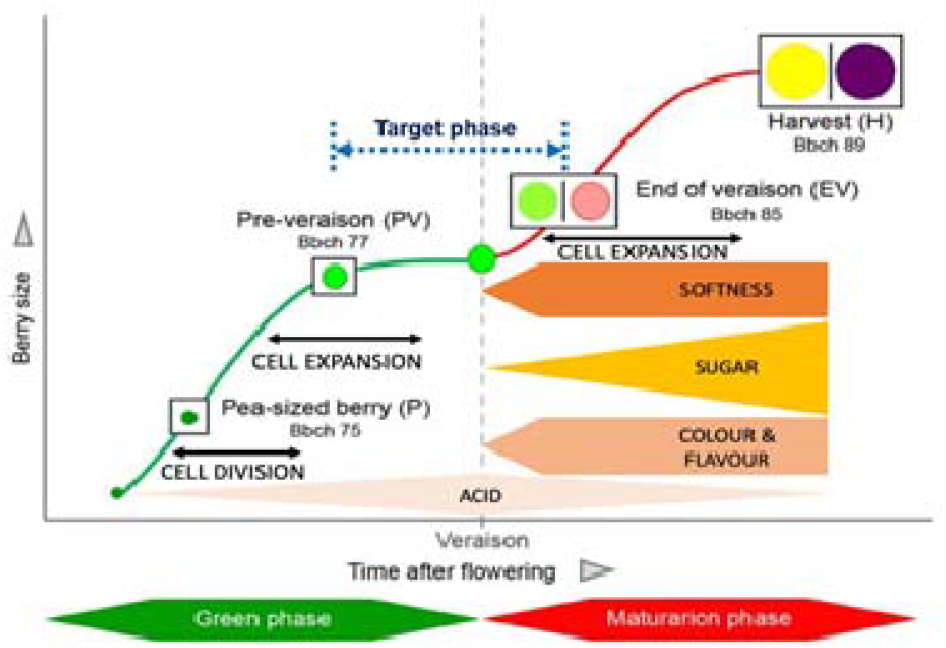
Schematic representation of berry grape development. Modified from Massonet al (2017) and Robinson and Davies (2000). The curve specifies changes in berry size. The inception of ripening (véraison) is shown by the vertical dash line. Softening, sugar accumulation and metabolism of organic acids and synthesis of colour and flavour compounds all occur after véraison.

The few eQTL studies in grapes have been carried out, mainly, to study a group of genes of the same pathway [24, 25] and in general studies that surveyed genome-wide expression to study ripening in grapes are scarce. These studies focused on specific pathways studied some candidate genes such as *VvMYBC2-L1*, a gene coding for an R2R3-MYBprotein and is involved in regulating PA synthesis, which are synthesized from anthesis to véraison. The VIT_16s0039g02230 gene (*VvUFGT*), which seems to be necessary in grapes for the transition of colourless anthocyanidins into their coloured glucosyl conjugates [31] or the VIT_02s0033g00410 gene (*VvMYBA1*) encoding an R2R3 MYB-type transcription factor [32]. Interestedly, from the 21 eQTLs identified by [24], 17 were novel loci that not correspond with known cis-regulatory sequences or candidate transcription factors. These results point out the importance to develop studies that surveyed genome-wide expression to identify an extended inventory of cis-regulatory sequences associated with grape ripening.

In this sense, the aim of this study is to characterize the genetics of the gene expression levels during ripening of fruit of *Vitis vinifera* L. through the use of bioinformatics approaches commonly used in human research. The final inventory of cis-eQTLs will be an important resource for future research to understand the mechanistic basis for variation in gene regulation during ripening in this species, and could be considered general markers of ripening in grapes.

## Materials and methods

### 2.1 Plant Material, RNA Extraction and Library Preparation and Sequencing

Five red-skinned grape cultivars (“Barbera nera”, “Sangiovese”, “Refosco”, “Negro amaro” and “Primitivo”) and five white-skinned grape cultivars (“Garganega”, “Vermentino”, “Moscato bianco”, “Glera”, and “Passerina”) were used in the current study. The 10 cultivars were grafted onto SO4 rootstock and grown in the same vineyard at the CREA-VE grapevine germplasm collection (Susegana, Veneto region, Italy). All vines received similar cultural practices (disease control, soil management, pruning, fertilization and irrigation) and were cultivated with the same training system and on homogenous soil and had the same age. Four different growth stages were considered for each variety: pea-sized berries (Bbch 75) at almost 20 days after flowering (P), berries beginning to touch (Bbch77) just prior véraison (PV or Touch), the softening of the berries (Bbch 85) at the end of véraison (EV or Soft) and berries ripe for harvest (Bbch 89) (H) (Figure 1). A total of 100 berries were collected from five vines of each variety block, at each stage of the development. Berries were collected from both sides of the canopy, and from different areas of the clusters. A more detailed description about berries collection can be found in [30].

### 2.2 DNA-seq Data Analysis

According to the published studies, genomic DNA from each cultivar was sonicated and paired-end libraries, 2 × 100-bp, were generated and (sample preparation guide, Illumina Inc., https://www.illumina.com) sequenced on Illumina HiSeq2500 sequencing machine. The raw data of “Vermentino” and “Sangiovese” were deposited in the National Center for Biotechnology Information (NCBI) Short Read Archive (SRA, https://www.ncbi.nlm.nih.gov/sra) under the BioProjects SRP106422 [33] and SRP111078 [25], respectively. The rest of the data for each cultivar used were obtained from BioProjects PRJNA373967, all sets of sequences were downloaded using a software module called the sratoolkit (Table S1).

The main difference regarding DNA data processing, with the previously published study of these cultivars is the use of a different pipeline for SNP calling. The general workflow used here can be observed in Figure 2. Briefly, in order to evaluate the general quality of reads in each file the FastQC package was used. The command used can take multiple files as input and outputs a quality report in html format. After that, low quality bases and/or adapter contamination were removed from reads using [34]. This step would discard any read trimmed shorter than 45bp. FastQC was run again on the trimmed data to confirm and ensure the final quality of reads. After this second quality control (QC) step, and before the alignment step, the reference genome was indexed using the command BWA index. Then, the V. vinifer*a* 12Xv2 reference genome [35] of the strain PN40024 (GCA_000003745.2) were used to align the reads using BWA [36]. BWA is one of the most widely used short-read aligners. BWA implements several alignment methods, but mem was selected. Reads were aligned with the default parameters. The output from an aligner such as BWA are in Sequence Alignment/Map (SAM) format. After this process the SAM files were compressed to Binary Alignment Map (BAM) format using samtools [37]. Reads were sorted by their positions in the reference genome using picard tools [38]. A BAM index was created and the variant calling was carried out using the methods implemented in bcftools. Finally, to remove low quality variants (e.g., variants with a low read depth or variants only supported by poorly aligned reads) from our data set, a quality filtering was applied to the VCF file using the command bcftools view (bcftools view -i ‘QUAL>19 && DP>2$ && (AC/AN)>0.05 && MQ>20’). In our study, the VCF file was used for linkage disequilibrium (LD) decay analysis, using the software PopLDdecay [39].

**Figure 2:**
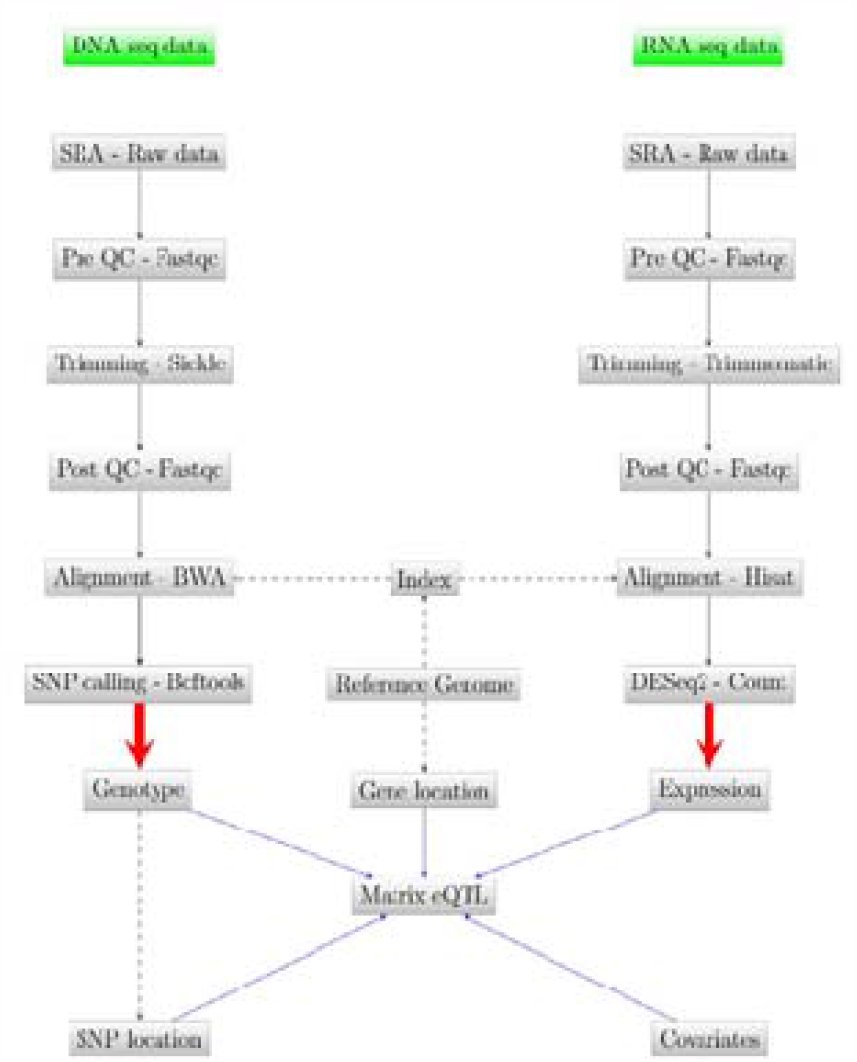
General workflow of the study.

### 2.3 RNA-seq Data Analysis

A total of 400mg of RNA was extracted from berry pericarp tissue (entire berries without seeds), a detailed description about RNA extraction and library preparation and sequencing of the 120 samples (10 varieties at four stages, in total 40 triplicate samples) can be also found in [30]. The complete information of the yielded 120 SRA files, downloaded from two BioProjects PRJNA265040 and PRJNA265039, can be found in Table S2 and Table S3. All the steps for RNA-seq analysis were performed in the UConn CBC Xanadu cluster belonging to the University of Connecticut, USA. This cluster also has SLURM as managing software. The general workflow used in this part can be observed in Figure 2. For RNA data, after the quality control (QC) step, Trimmomatic [40] was used to trim low quality and adapter contaminated sequences. In this case, the alignment of reads to the reference genome was performed by HISAT2 [41]. HISAT2 is a fast and sensitive aligner for mapping next generation sequencing reads against a reference genome. Before the alignment, the hisat2build module was used to make a HISAT index file for the genome. By default, HISAT2 outputs the alignments in SAM format. Again samtools was used to sort the sequences, convert them to binary format and compress them. Finally, the function htseq-count from the HTSeq python package[42] was used to count how many reads map to each annotated exon (gene) in the genome. The final count for each gene was obtained from sum values for all their exons. These final counts per gene are the inputs of the R package DESeq2 [43], used for the differential expression analysis.

### 2.4 Matrix eQTL

Matrix eQTL [44] was used for the eQTL analysis. This software was created to handle expression and large genotype data sets. Matrix eQTL checks for association between each SNP and each transcript by modeling the effect of genotype as either categorical (ANOVA model) or additive linear (least squares model). The simple linear regression (used in this study) is one of the most commonly used models for eQTL analysis. In addition, Matrix eQTL can test for the significance of genotype-covariate interaction (not con¬sidered in this study). Matrix eQTL also supports correlated errors to account for relatedness of the samples and heteroscedastic. Matrix eQTL implements a separate test for each gene-SNP pair and corrects for multiple comparisons by calculating false discovery rate (FDR) [45]. Five different input files (snps=snps data; gene=expression data; cvrt=covariates; genepos = gene location; snpspos = SNP location) are required to run Matrix eQTL All these files need to have a specific format. The columns of all three files must have matching order, corresponding in each column to a sample and with one gene/SNP/covariate in each row. In the case of the genotype file, if a linear model is used, as in this study, the values must be numerical in this data set. For that reason, extract.gt function from the R package vcfR [46] was used to read and extract genotypes, from our VCF filtered file, in numeric format. The p-value threshold for cis-eQTLs (pvOutputThreshold.cis) in this study was 1e-8 and the maximum distance at which gene-SNP pair is considered local (cisDist) was 1000. In our study, covariates were not considered. The location of each gene was obtained from the gff3 file: Vitis_vinifera_gene_annotation_on_V2_20.gff3 available at https://urgi.versailles.inra.fr/Species/Vitis/Annotations. The location of each SNP was obtained from the VCF file obtained.

Matrix eQTL was run six times:

1. To detect cis eQTLs in white cultivars for Touch stage (TW)
2. To detect cis eQTLs in white cultivars for Soft stage (SW)
3. To detect cis eQTLs in red cultivars for Touch stage (TR)
4. To detect cis eQTLs in red cultivars for Soft stage (SR)
5. A general analysis to detect cis eQTLs for Soft stage (SG)
6. A general analysis to detect cis eQTLs for Touch stage (TG)

As a result of each run, each significant gene-SNP association was recorded in a separate line in the output file. Every record contains a SNP name, a transcript name, estimate of the effect size, tor F-statistic, p-value, and FDR.

Comparison of detected cis eQTLs between samples After Matrix eQTL, several comparisons were performed to detect intersection(s) of significant gene-SNP associations between samples using a ven-n/Euler diagram (http://bioinformatics.psb.ugent.be/webtools/Venn/). It is a quick approach to indicate which elements are in each intersection or are unique to certain conditions. The comparisons were:

1. Soft White (SW) vs Touch White (TW)
2. Soft Red (SR) vs Touch Red (TR)
3. Touch Red (TR) vs Touch White (TW)
4. Soft Red (SR) vs Soft White (SW)
5. General Touch (TG) vs General Soft (SG)

Prediction of genetics variants effects was performed using SnpEff v4.3e [47] based on the grape gene annotation (Vitis_vinifera_gene_annotation_on_V2_20.gff3; https://urgi.versailles.inra.fr/Species/Vitis/Annotations). The SNP predicted effects were categorized by impact, as moderate (non-synonymous substitution); modifier (with impact on noncoding regions); low (synonymous substitution); or high (disruptive impact on the protein).

### 2.5 Enrimencht Analysis and protein annotation

Using the list of unique and intersect IDs of each sample condition (e.g. Soft White, Touch White, Soft Red, Touch Red, etc.) in AgriGO version 2.0 a Singular Enrichment Analysis (SEA) [48] was performed. This traditional approach, SEA, iteratively evaluates individual annotation terms one at a time against an established list of candidate genes for enrichment. After comparing the observed frequency of an annotation term with the frequency expected by chance an enrichment p-value is estimated; individual terms are deemed enriched if they past some cut-off (e.g. p-value ≤ 0.05) [49]. In the present study, Fisher method was used for the statistical test. Actually AgriGO uses version1 of the grape genome transcript (genoscope) so each term was converted from VCost.v3 to V1 using the file list_genes_vitis_correspondencesV3_1.xlsx available at https://urgi.versailles.inra.fr/Species/Vitis/Annotations. Protein annotation was carried out using UniprotKB [50] and DAVID [51].

## Results

### 3.1 DNA data analysis

Basic statistics of the alignment of the trimmed sequences to the reference genome of each replicate and cultivar can be found in Table S4 and Table S5. The summary of each cultivar are shown in Table 1. This information was obtained through the command samtools stats. From the total number 2,741,939,082, 93% of the reads (2,550,198,134) were mapped with a global average quality (ratio between sum of bases qualities and total length) of 37.05. The percentage of reads properly paired out was 81.3%. The cultivars with the lower number of total and mapped sequences were “Vermentino” and “Primitivo”. The lower value of average quality was observed in “Sangiovese” (35.4) and “Vermentino” showed the lowest percentage of properly paired reads (62.8).

**Table 1:** Statistics of the alignment of reads by bwa

| Cultivar | Raw total sequences | Filtered sequences | Reads mapped | Average quality | Percentage of properly paired reads (%) |
| --- | --- | --- | --- | --- | --- |
| Barberanera | 579,559,276 | 0 | 528,667,032 | 37.4 | 75.1 |
| Primitivo | 82,920,496 | 0 | 69,815,801 | 37.2 | 71.1 |
| NegroAmaro | 106,016,336 | 0 | 96,165,208 | 36.7 | 66.2 |
| Vermentino | 69,361,256 | 0 | 65,617,979 | 36.4 | 62.8 |
| Glera | 288,557,208 | 0 | 271,496,346 | 37 | 83.9 |
| Garganega | 259,300,044 | 0 | 242,322,498 | 37 | 80.7 |
| Refosco | 164,876,640 | 0 | 152,540,110 | 38.2 | 81.9 |
| Sangiovese | 589,391,258 | 0 | 556,212,326 | 35.4 | 82.4 |
| Moscato Bianco | 309,292,648 | 0 | 292,442,700 | 37.1 | 83.9 |
| Passerina | 292,663,920 | 0 | 274,918,134 | 37.1 | 82.4 |
| Total | 2,741,939,082 | 0 | 2,550,198,134 | 37.05 | 81.3 |

In total 17,282,868 genetic variants were obtained after running the pipeline used in this study. A total of 560,417 small insertions, 519,025 deletions (InDels) and a total 16,203,426 SNPs were detected. This set was filtered to remove SNPs with two (or three) possible alternative alleles in the VCF, obtaining a total of 15,692,912 SNPs. This set was used to calculate the number of transitions and transversions. In this final set, 70% were transitions (11,111,319 loci) and 29.2% transversions (4,581,593 loci), and the ratio of transition/transversion was 2.43. A/G and C/T transitions occurred at the highest and very similar frequencies. The four different types of transversions presented different frequencies 10.053% for A/T, 7.47% for A/C, 4.22% for C/G and 7.46% for G/T (Table 2).

**Table 2:** Classification of SNPs based on their nucleotide substitutions, either transitions (Ts) or transversions (Tv).

|  | Transitions |  | Transversions |  |  |  |
| --- | --- | --- | --- | --- | --- | --- |
|  | AG | CT | AC | AT | CG | GT |
| Number of allelic sites | 5,551,580 | 5,559,739 | 1,172,883 | 1,576,561 | 662,107 | 1,170,042 |
| Percentage of allelic sites | 35.38 | 35.43 | 7.47 | 10.05 | 4.22 | 7.46 |
| Total(Percentage) | 11,111,319 (70.80%) |  | 4,581,593 (29.20%) |  |  |  |

Additional filtering to remove sites with missing data and also homozygous sites were performed using a custom script in python. As a result, 12,198,767 SNPs were retained for red cultivars and 11,128,404 SNPs for white cultivars. For the general analysis (using all samples) a total of 11,895,933 SNPs were retained.

### 3.2 LD analysis

LD is a very important effect that plays a big role in genetic association studies. It describes the effect that genetic variants are not always inherited independently due to recombination atterns during reproduction. A high LD, confirm that SNPs associated with a gene are highly linked to each other and therefore describe the same effect. The results obtained by PopLDdecay showed that linkage disequilibrium decay to 0.2 around 2kb and to 0.1 around 300kb (Figure 3).

**Figure 3:**
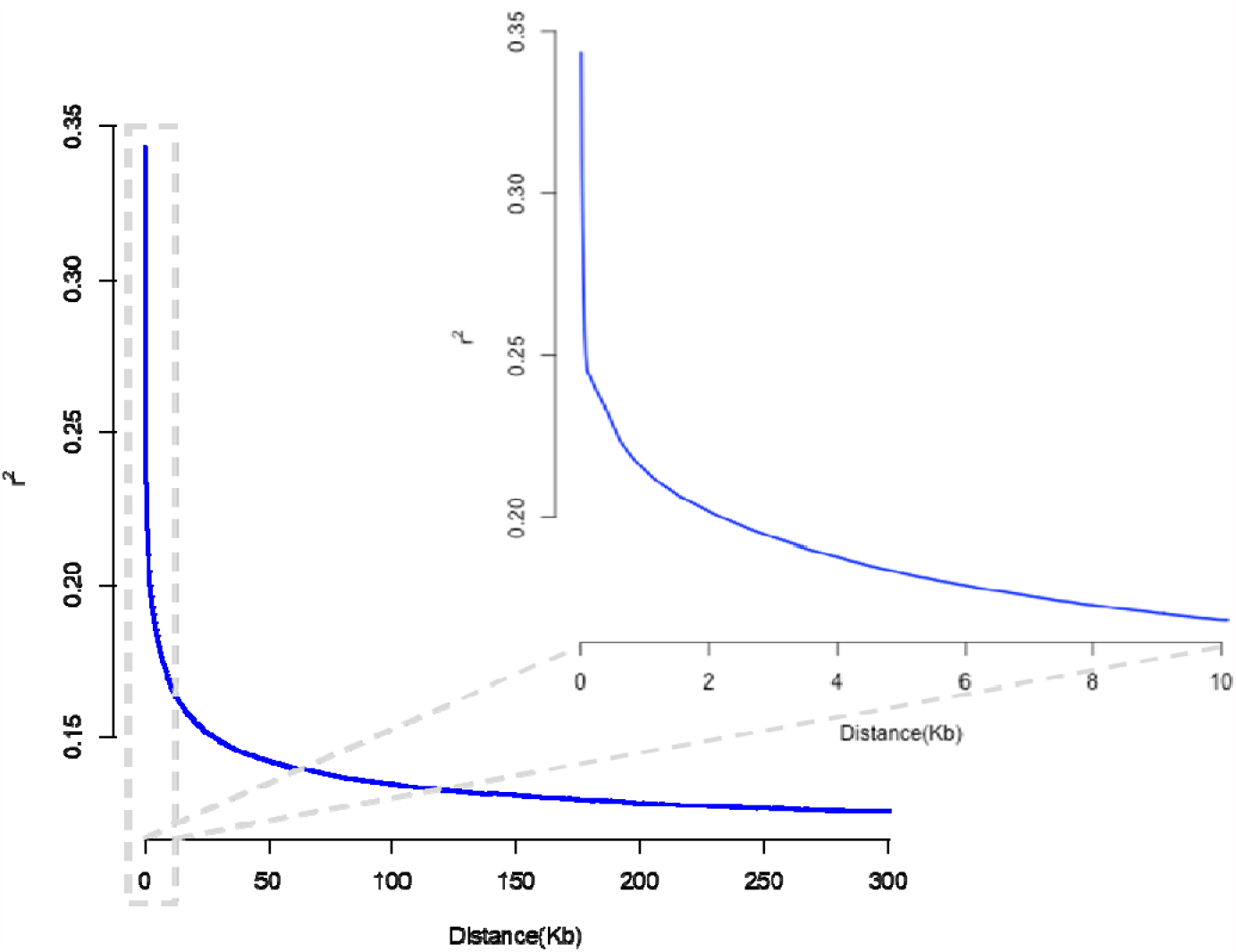
Plot of LD decay.

### 3.3 Expression data analysis

The detection of the differential expression genes (DEGs) between the selected conditions (soft and touch) in each type of cultivar (white and red), and in general, are needed for downstream analysis. The complete and exhaustive pipeline was carried to process a large amount of expression data. For red cultivars, a total of 208,14 Gbases and 139,03 Gbytes of data were obtained, representing a total of 2,174,234,253 sequences (Table S4). The total trimmed sequences were 1,905,562,540 (87,64% trimmed), the total number of lost sequences was 268671713, a total of 1,692,026,464 sequences were aligned (88,79% of the total), being 213,536,076 not aligned (11,21% of the total). From this set, 44,436,905 were aligned more than one (2,33% del total) and 1,647,589,559 were aligned just one time (86,46%) (Table S4). In the case of white cultivars, a total of 2,163,260,617 sequences (Table S5) were used representing a total of 207,23 Gbases and 138,65 Gbytes of data (Table S3). The total trimmed sequences was 1,911,460,644 (88,36% trimmed), the total number of lost sequences was 251,799,973, a total of 1,702,609,789 sequences were aligned (88,36% of the total), being 208,850,855 sequences not aligned to the reference (10,93% of the total). From this set, 46,507,687 were aligned more than one (2,43% del total) and 1,656,102,102 were aligned just one time (86,64%) (Table S5). The data quality was assessed in parallel to the differential expression analysis, this essential step was performed with the PCA function. The goal here is to detect the data points obtained, from a specific samples or several samples in which experimental treatment suffered from an abnormality, which can clearly affect to our study. The PCA plots show the samples in each comparison in the 2D plane spanned by their first two principal components (Figure 4). In all three comparisons PCA1 explained around 70%, means that the growth stage has a larger effect and that in general, replicates from each sample showed low variability between them, which can ensure the absence of a well-known batch effect, something not considered in this study.

**Figure 4:**
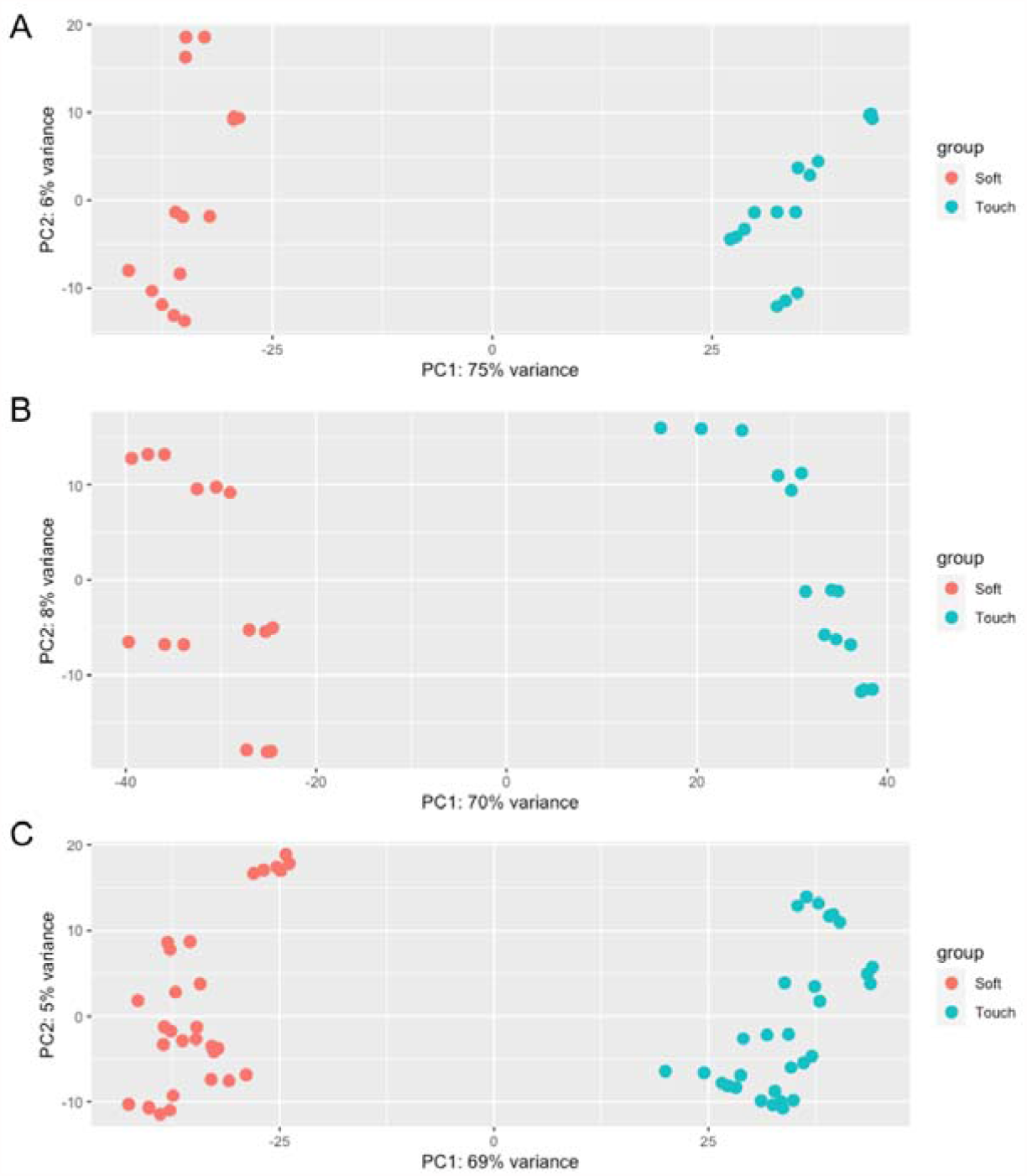
Principal Component Analysis between soft and touch conditions. A) Red grapes B) White grapes c) General analysis.

The total number of annotated genes in the grape genome is 42,416 (Vitis_vinifera_gene_annotation_on_V2_20.gff3). In the case of red cultivars, a total of 25,470 (60%) genes with nonzero total read count were detected, after filtering out genes with counts <20 and adjusted p-value cutoff (FDR) >0.05, 19,379 genes were retained. In this final set, the total number of genes with a LFC (log2 foldchanges) > 0 (up) was 5,903 (30%) and with a LFC < 0 (down) was 5996 (31%). In the case of white cultivars, a total of 25,087 (59%) genes had nonzero total read count. From this set, after filtering by counts and FDR, 19,160 genes were retained. The total number of genes with a LFC (log2 foldchanges) > 0 was 5,554 (29%) and with a LFC < 0 (down) was 5,280 (28%). In the case of the general study, 26,691 had nonzero read counts and after applying both filters, 20621 genes were retained, with 6,641 (32%) up and 6,806 (33%) down regulated.

### 3.4 Matrix eQTL

As results of Matrix eQTL in total for all comparisons 525 genes were identified, representing 2.05% of the total number of genes. The number of associated genetic variants was 10,618 (7,811 unique) (Table 3). The SR condition showed the highest number of genes, with a total of 175 genes detected.

**Table 3.** Number of genes and SNPs per each condition analyzed.

| Condition | Genes | SNPs |
| --- | --- | --- |
| General_SOFT | 141 | 1415 |
| General_TOUCH | 110 | 753 |
| Red_SOFT | 175 | 2822 |
| Red_TOUCH | 96 | 1475 |
| White_SOFT | 159 | 2857 |
| White_TOUCH | 66 | 1296 |
| **Total** | **747** | **10618** |

In this case, the number of SNPs per gene ranged from 1 SNP to 213 SNPs (Vitvi19g00958). In the case of TR a total 96 genes were detected with 1 SNP to 188 SNPs (Vitvi18g01761). The results in the case of white cultivars showed 159 genes for SW and 66 for TW. The number of SNPs per gene was also variable in the white cultivars, oscillating from 1 SNP to 192 SNPs (Vitvi13g01155) per gene, in both cases. The general analysis for soft (SG) and touch (TG) showed a total of 141 and 110 genes, respectively. The number of SNPs per gene varied from 1 SNP to 317 SNPs (Vitvi11g01142) in the case of soft and from 1 to 64 SNPs (Vitvi08g02005) per gene in the case of touch. The complete information about gene ID and SNPs by each comparison can be found in Table S6.

The Venn diagram analysis, in the comparison SW vs TW, showed a total of 33 genes shared between both (Figure 5A). A total of 126 genes were unique to SW and 33 to TW. The comparison between SR and TR, showed a total of 44 shared genes between both conditions (Figure 5B). The SR showed 131 unique elements and 53 genes were unique for TR. The comparison between TR and TW, showed that only 6 genes were shared between these conditions (Figure 5C). The TR showed 91 unique elements and 60 genes were unique to TW. In the comparison SW vs SR, the venn diagram showed that 27 genes were shared between both (Figure 5D). A set of 132 genes were unique to SW and 148 genes unique to SR. The TG vs SG comparison showed that 62 elements were unique to TG, 93 were unique to SG and 48 elements were shared between both conditions (Figure 5E).

**Figure 5:**
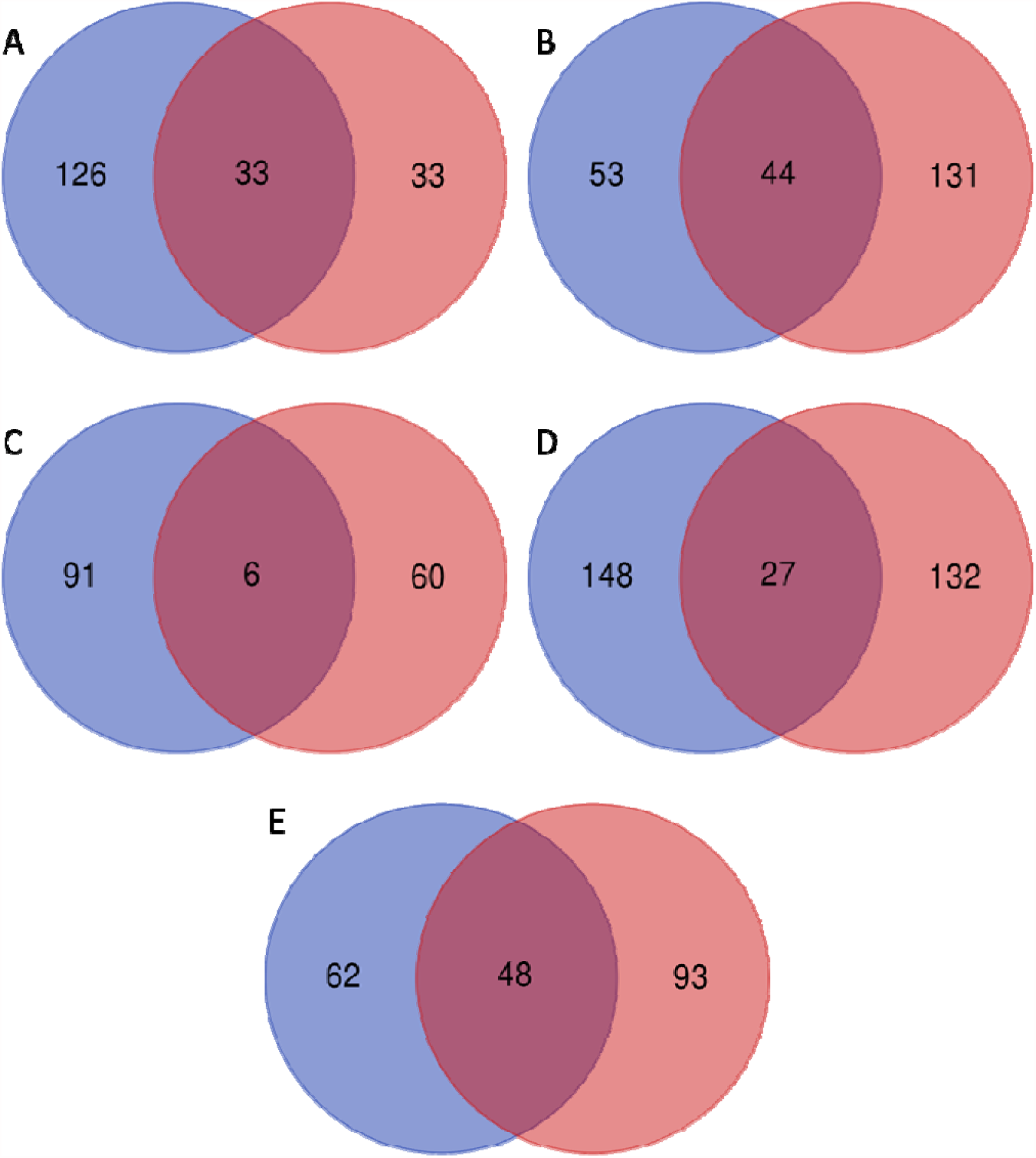
Venn diagram for each comparison. A)SWvsTW; B)SRvsTR; C)TRvsTW; D)SWvsSR; E)TGvsSG.

### 3.5 Functional classification of DEGs

The individual terms detected by SEA beyond the determined cut-off (p-value ≤ 0.05) were deemed enriched. The complete list of terms can be found in Table S7. The number of terms was variable for each comparison. The number of GO terms associated to each gene and each condition can be found in Table S8. In the case of DEGs in SW (159=126+33) after comparing SW and TW, the SEA analysis using the genome reference, highlighted three terms. Electron carrier activity (GO:0009055; p-value 0.00017, FDR 0.031) in the molecular function (MF) category, with 5 genes (3.77%) associated and intracellular membrane-bounded organelle (GO:0043231; p-value 0.00016, FDR 0.0041) and membrane-bounded organelle (GO:0043227; p-value 0.00016, FDR 0.0041) in the cellular components (CC) category, with 13 genes (8.17%) associated in each category. No GO term was significantly enriched among DEGs in TW (66=33+33).

After the comparison of TR vs SR, the SEA of the elements (97=53+44) observed in TR showed six significant GO terms. Five of them, Metal ion binding (GO:0046872; p-value 3e-06), cation binding (GO:0043169; p-value 3.2e-06), ion binding (GO:0043167; p-value 7.3e-06), transition metal ion binding (GO:0046914; p-value 3.8e-05) and zinc ion binding (GO:0008270; p-value 0.0012), in the molecular function category, with 16 (16.49%), 16 (16.49%),16 (16.49%), 12 (12.37%), 7 (7,22%) genes associated. The other term was single-organism process (GO:0044699; p-value 0.00033) in the biological process category to which 20 genes (20,62%) were associated. In the case of the list of elements of SR (175=131+44), the SEA showed a total of 23 significant GO terms. The most significant GO terms were multicellular organism development, (GO:0007275; p-value 5.7e-06), single-organism developmental process, (GO:0044767; p-value 8.1e-06), single-multicellular organism process (GO:0044707 p-value 5.7e-06) and anatomical structure development (GO:0048856 p-value 8.1e-06) with very low FDR (0.00086), all of them in the Biological Process category and with a total of 6 genes (3.42%) associated to each term.

After the comparison of TR vs TW, the SEA showed that in the case of TW (66=60+6) no significant GO terms were detected. In the case of the list of elements of TR (97=91+6), the significant GO terms were the same identified previously in TR in the comparison TR vs SR. The SEA of the list of elements of SR (175=148+27) after comparing SR vs SW showed the same 23 GO terms identified previously for SR in the comparison TR vs SR. In the case of SW (159=132+27) the same three significant GO terms, as in the case of SW in the comparison SW vs TW, were identified.

After the general comparison TG vs SG, the SEA for the TG list (110=62+48) showed 13 significant GO terms, all described in the Biological Process category. The most significant term response to abiotic stimulus (GO:0009628;p-value 6.60E-09, FDR 1.70E-06) to which 5 genes (4.55%) were associated (Figure S1). In the SG (141=93+48), 15 significant GO terms were 1identified, being the most significant term defense response (GO:0006952; p-value 8.3e-09, FDR 3.1E-06) to which 7 genes (4.96%) were associated (Figure S2). In summary, the SEA detected a short list of 39 significant GO Terms between these both stages of development in all comparisons, representing a unique list of 114 associated genes.

### 3.6 SNP effect annotation

To explore the effects of the total variants (7811) on these genes, the software SnpEff detected a total of 20,219 effects, 0.89% (181) were high (with disruptive impact on the protein), 2.50% (506 effects) were low effect (with synonymous substitution), 4.32% (875) were moderate (genetic variation could have a non-synonymous substitution) and 92.27% (18,657) were modifier effects (with impact on noncoding regions) (Figure 6). The most common functional class of the effect was missense with 902 effects (65.9%), silent class was observed in 28% of the effects (Figure 7). The ratio missense/silent was 2.28%.

**Figure 6:**
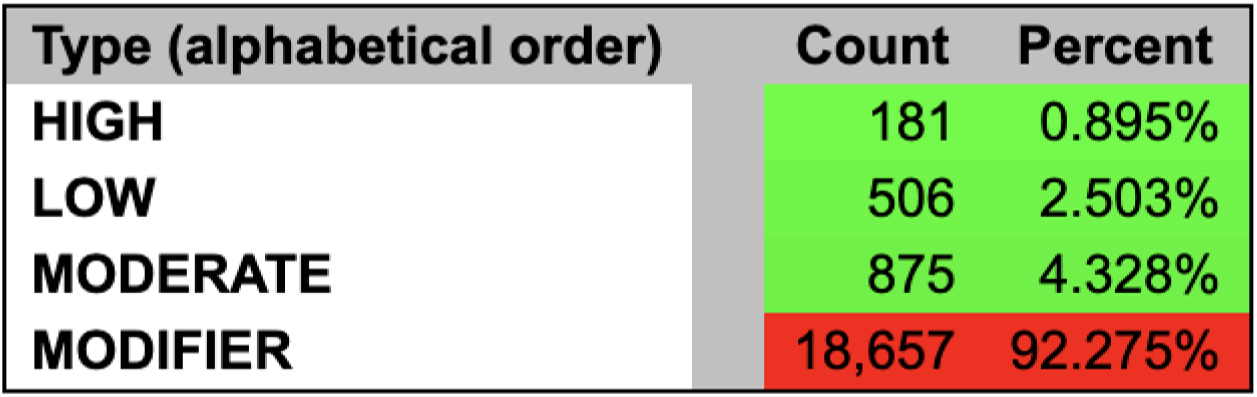
Number of effects by impact.

**Figure 7:**
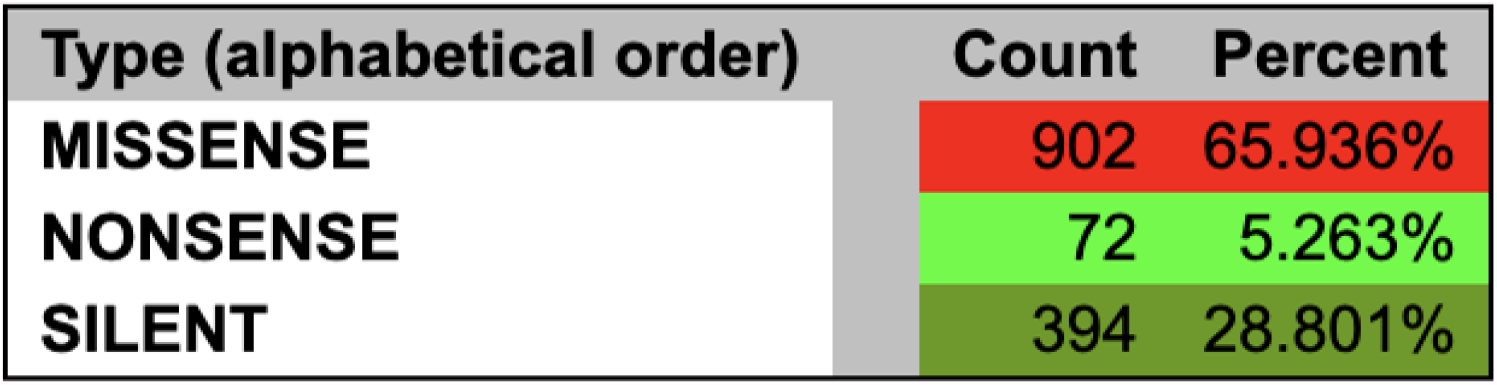
Number of effects by functional class.

In the category of SNPs with a modifier effect, a default length of 5 kbp downstream of the most distal polyA addition site was the most frequent variant (27.08%) (Supplemental file 1). The most frequent mutations for SNPs with a low effect were splice region variants (0,565%), while missense variants (4,213%) within SNPs with a moderate effect, were the most frequent mutation. Finally, SNPs with a high impact had stop gained variants and frameshift variants as the most frequent mutations (0,379% and 0,241%, respectively). The majority of effects were located in downstream and intron regions with 27.24% and 23.60%, respectively, being the splice site donor the region with the lowest percentage of effects 0.025% (Figure 8).

**Figure 8:** Distribution of variants across genomic regions.

### 3.7 Protein annotation

To obtain the functional information of the 525 candidate genes associated with eQTLs, UniProtKB and DAVID were used. A 70% of them, 367, received a UniProtKB identifiers, and 44 (8.39%) received a DAVID term. The annotation information for each gene can be found in Table S9. The five genes associated with GO:0009055 in the SW list (159) coding for uncharacterized proteins. In the set of 13 genes associated with GO:0043231 and GO:0043227, two of them coding for uncharacterized proteins and the rest coding for proteins such as growth-regulating factor, Phospholipase D (EC 3.1.4.4), CBFD_NFYB_HMF domain-containing protein, Homeobox domain-containing protein, 3-hydroxy-3-methylglutarate-CoA lyase (EC 4.1.3.4) (Hydroxymethylglutaryl-CoA lyase, mitochondrial), peptidase_S9 domain-containing protein, ATP-dependent Clp protease proteolytic subunit (EC 3.4.21.92), AP2/ERF domain-containing protein, PMR5N domain-containing protein, auxin-responsive protein, MADS-box domain-containing protein.

From the total 16 genes of the TR lists (97=53+44) associated with five significant GO terms (Metal ion binding (GO:004687), cation binding (GO:0043169), ion binding (GO:0043167), transition metal ion binding (GO:0046914) and zinc ion binding (GO:0008270)), in the molecular function category, eight genes coding for uncharacterized proteins, the rest coding for proteins such as RING-type domain-containing protein, PHD finger protein ING, HMA domain-containing protein, PKS_ER domain-containing protein, Germin-like protein, S-acyltransferase (EC 2.3.1.225) (Palmitoyltransferase) or Phytocyanin domain-containing protein. The 20 genes associated with the other significant term (single-organism process (GO:0044699)) in the biological process category, coding for proteins such as Kinesin motor domain-containing protein, FAD-binding PCMH-type domaincontaining protein, PKS_ER domain-containing protein, TIR domain-containing protein, Thiaminephosphate pyrophosphorylase (EC 2.5.1.3), Phospholipase D (EC 3.1.4.4), (Thiamine-phosphate synthase), Non-specific lipid-transfer protein, S-acyltransferase (EC 2.3.1.225) (Palmitoyltransferase), DLH domain-containing protein.

In the case of the list of elements of SR (175=131+44), the six genes associated with the most significant GO terms (multicellular organism development (GO:0007275), single-organism developmental process (GO:0044767), single-multicellular organism process (GO:0044707) and anatomical structure development (GO:0048856)) coding for several proteins such as GID1b, SANT domain-containing protein, eukaryotic translation initiation factor 3 subunit H (eIF3h) and DLH domain-containing protein.

In the TG list (110=62+48), four of the five genes associated with the most significant term response to abiotic stimulus (GO:0009628) coding for an uncharacterized protein, and only one gene (Vitvi04g00189) coding for a known protein, SAM domain-containing protein. In the SG (141=93+48) list, the seven genes associated with the significant term defense response (GO:0006952) coding for proteins such as AAA domain-containing protein, TIR domain-containing protein and ANK_REP_REGION domain-containing protein.

### 3.8 Description of associated genetic variants with gene expression

The group of 181 genetic variants with potential to have a high impact on gene function identified in this study, are affecting 110 genes in the grape genome (Table S11). In the majority of the genes at least one SNP, two or three SNPs were observed, except in Vitvi06g00918 coding for a Homeobox domain-containing protein, Vitvi12g01425 (unknown), Vitvi16g00122 coding for a PNP_UDP_1 domain-containing protein, with four, Vitvi16g01859 coding for a putative F-box protein At5g55150 (LOC104882163) with six, Vitvi18g01761 (Uncharacterized protein) with seven and Vitvi05g02226, coding for a putative disease resistance RPP13-like protein 2(LOC100852873), with 13 (Table S11). One candidate gene with one high impact genetic variant according to snpEff was Vitvi06g00964 (VIT_06s0009g00820). This gene coding for a Kinesin motor domain-containing protein and was only found in red cultivars in both conditions and in both general analysis. A total of 51 genetic variants affecting the gene expression of this gene were detected by Matrix eQTL in the general analysis of touch and soft and 76 genetic variants in the red soft analysis. All the detected eQTLs, in the case of red cultivars or white cultivars, separately, showed only two genotypes categories (AA and AB) or (BB and AB). Only in the general comparison, using all cultivars (red+white), 77 eQTLs showed three genotypes categories in the soft comparison and 31 eQTL in the case of touch condition showed the three genotypes (AA, AB, BB). This total number of eQTLs were associated with only 14 genes (Table S10). In the case of Vitvi05g01272, a change from CC to TT supposed more than 900% of increment of the expression (Figure 9) and in the case of Vitvi15g01618 a change from GG to AA supposed an increment of 500% in the expression of the gene.

**Figure 9:**
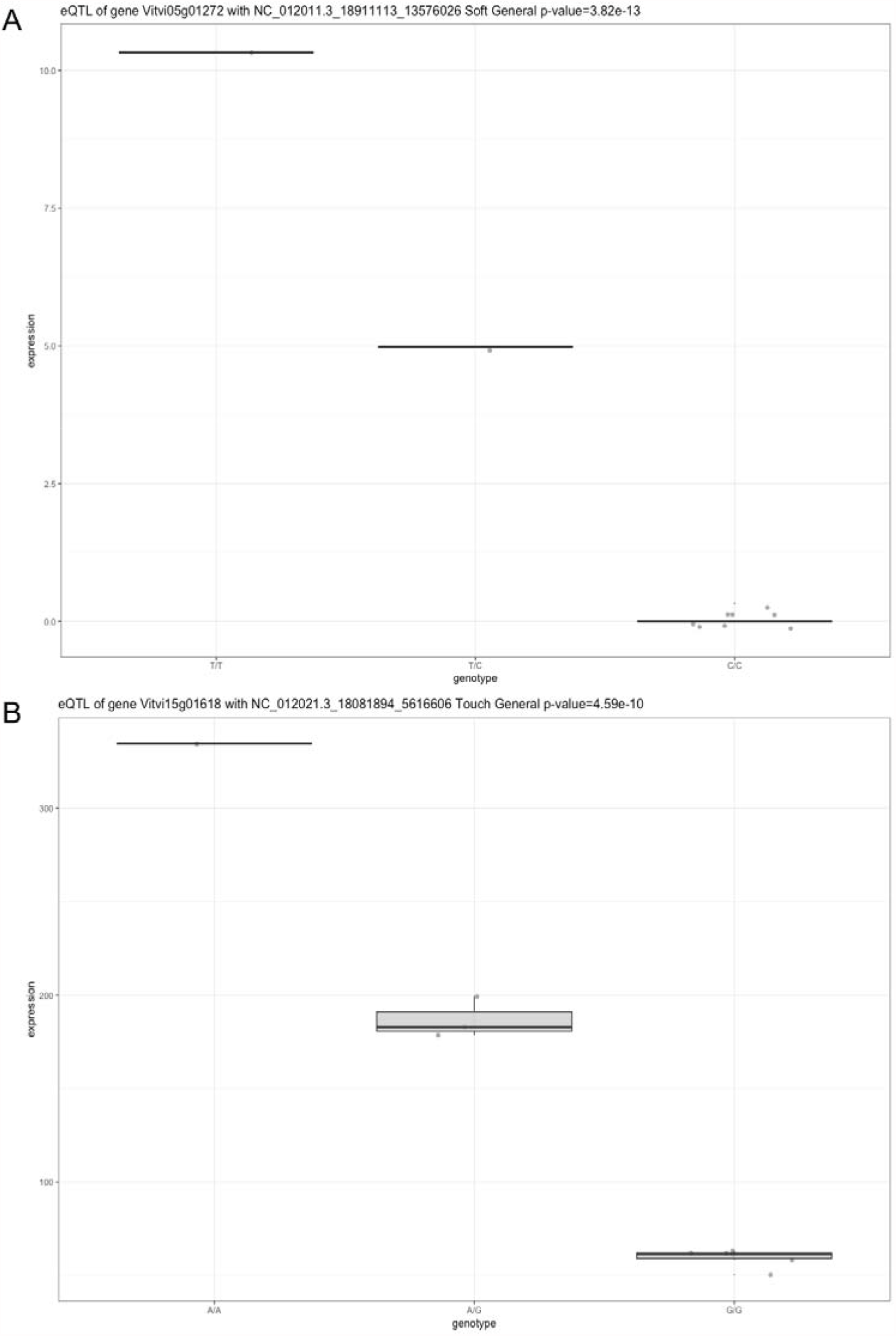
Differential expression of genes associated with a eQTL. A)Vitvi05g0127. B)Vitvi15g01618

In the list of genes with no HIGH impact variants obtained by snpEff is Vitvi09g00107. This gene coding for growth-regulating factor was found only in white cultivars in the soft stage by Matrix eQTL with five genetic variants affecting to the expression of this gene (NC_012015.3_1133980, NC_012015.3_1133985, NC_012015.3_1133986, NC_012015.3_1137239, NC_012015.3_1137290). The NC_012015.3_1133980 genetic variant was annotated as an INDEL in chromosome 9 of vitis vinifera, and with a MODIFIER effect. Similarly, the gene Vitvi11g00497 had no HIGH impact variant associated with its expression. This gene was only detected in soft stages in white and red cultivars and in the general soft analysis by Matrix eQTL. Eight different genetic variants (NC_012017.3_5097457, NC_012017.3_5097792_896739, NC_012017.3_5095382, NC_012017.3_5095554, NC_012017.3_5096958, NC_012017.3_5097909, NC_012017.3_5098320, NC_012017.3_5098472) were associated with changes of expression of this gene. The genetic variant NC_012017.3_5097457 (A/G) was detected in the three run of Matrix eQTL, and annotated with several LOW (5 prime UTR premature start codon gain variant) and MODIFIER (upstream gene variant) effects by snpEff. Another gene that was only detected in red cultivars in the touch stage was Vitvi14g02553, this gene coding for a Germin-like proteins (GLP). Four genetic variants were annotated by Matrix eQTL for this gene (NC_012020.3_2975100, NC_012020.3_2975313, NC_012020.3_2975857, NC_012020.3_2976143).

## Discussion and conclusions

A detailed bioinformatic pipeline has been established to study an important phenomenon such as fruit ripening in an important woody crop species such as *Vitis vinifera* L. The information about genetics of gene expression obtained in this study provides researchers with new scenarios to understand this genetically programmed process that is physiologically and biochemically irreversible. The two main criticisms of this study can be the low number of genotypes used, and that a high percentage (67,30%) of the identified genes coding for an uncharacterized protein. Both issues could be understood from the difficulty to work with woody crops and from the point of view that *Vitis vinifera* is not a real model species in plants as can be *Arabidopsis thaliana*. As commented by [52], in higher eukaryotes gene annotation projects present high complexity reflecting the complexity that survives in eukaryotic cells, and more important at the present time all of our genebuilds (representations of the transcriptome that exists in nature) are incomplete because scientists do not fully understand the transcriptome,

Our results have shown a fast decay of LD for this species. According to literature, thousands of years of widespread vegetative propagation has resulted in a weak domestication bottleneck in grages, which is consistent with a fast decay of linkage disequilibrium (LD) normally observed in this species [53], reaching values of r2=0.2 within a few Kb at most [54]. These high LD values can make it difficult to select at most one SNP per gene being associated with altered gene expression. The existence of many SNPs associated with a gene it is most likely that those SNPs are “in high LD”, and probably highly linked to each other and therefore they describe the same effect. The fast decay LD observed here could suggest that the genes detected could be regulated by different SNPs independently. In fact, different values of LD were observed across four genomic regions between wine eastern cultivars, wine western varieties, eastern table grapes and wild grapevine individuals by [55]. A fine mapping approach, using software as FINEMAP [56], or TreeMap [57], could be necessary to detect the lead eQTL signals on specific regions (and genes). Fine mapping has been carried out in plants as maize, to understand one of the key steps in its domestication [58].

### 4.1.1 Functional and genes characterization of both growth stage

In grapes, a comprehensive transcriptional and metabolic analysis was performed over two seasons to understand the dynamics of ripening of this nonclimacteric fruit. All the molecular and biochemical phases leading to the initiation of ripening harbor a level of complexity not fully understood yet [59]. The results reported by [59], identified several functional categories such as “development”, “metabolism”, “diverse/miscellanenous functions”, “cellular process”, “regulation overview” or “response to stimulus, stress” together with several not previously identified genes in the context of grape ripening. The same authors observed that abiotic (e.g. osmotic, temperature) stress responses and biotic (e.g. pathogens) increase during grape ripening [59]. More recently, [60] studied two grape genotypes “Nebbiolo” and “Barbera” and observed that during ripening the functional categories that most differentially changed between both cultivars were response to stress and secondary metabolism. The biological networks of secondary metabolism in grapevine, at two levels, regulatory and catalytic, was previously studied [61]. These authors performed an integrated approach, combining microarray analysis and gas chromatography-mass spectrometry analysis, skins of berries were sampled from pre-véraison to over-ripening, at five selected developmental stages. The evolution of aroma during ripening was studied through the comparison of these five stages. As results from the comparison between stage1 (31 E-L stage 75 BBCH) and stage 3 (36 E-L), which could be the nearest stages to the one used here, showed a large list of downregulated candidates genes such as VIT_16s0050g01150 (Heat shock protein 90-1), VIT_13s0019g00860 (Small heat-shock protein HSP17.5 Cytosolic class I), VIT_00s0463g00020 (Scarecrow transcription factor 5 (SCL5)), VIT_08s0007g05880 (Dehydration-induced protein (ERD15) (ABA signaling)), VIT_08s0007g05790 (Calmodulin) or VIT_15s0046g01440 (BZip transcription factor G-box binding factor 3) and also upregulated genes such as VIT_12s0028g03860 (Zinc finger (C3HC4-type ring finger) protein (RMA1)), VIT_10s0003g03190 (RNA recognition motif (RRM)-containing), VIT_09s0054g01780 (P300/CBP acetyltransferase-related protein 2 gene), VIT_08s0007g08160 (Telomere repeat binding factor Like TRFL10). In the present study, any of these candidate genes have been detected, but on the other hand, some of these functional categories detected by [59] such as “response to stimulus, stress” or “developmental process” were also identified in this study. All these results, can suggest that the genetic architecture underlying grape ripening is still far from being complete.

With the GO analysis obtained here, clear differences can be observed between the two stages of growth studied, in general the touch stage is characterized by response to abiotic stimulus and the red stage is characterized as a defense response. In this sense, exogenous stimuli (temperature, light, abscisic acid, jasmonic acid and oxidative stress,) have a lower impact in véraison-stage berries, which seem to have a greater resilience to metabolic alteration driven by these factors, than pre-véraisonstage berries [62]. Probably, the accumulation of polyphenols in early stages, offer a strategy for the defense of the ripening berry [63]; otherwise, the stress input is overcome by the developmentally regulated metabolic program. More interestingly, a large amount of gene IDs identified by Matrix eQTL have not previously been identified in the context of grape ripening. The obtained characterization of regulatory variants can now provide a valuable resource to help the biological understanding of grape ripening.

### 4.1.2 Auxins and other candidates genes

According to the literature, after véraison stage, the content of several hormones, as auxins and cytokinins, decrease while abscisic acid concentration increases [64]. Auxins are phytohormones associated with an extensive variety of phases of the development including responses to various stimuli and fruit development. In addition, of the classical link with growth of these phytohormones at early stages of fruit development, others functions, such as the ability to affect ripening, have been inferred [65]. Recently, the study of their functions in the control of grape berry ripening has been tackled for several authors. Some of these studies have shown that in flowers and young berries concentrations of indole-3-acetic acid (IAA) are high, but rapidly decrease and prior to veraison (the initiation of ripening) their concentrations are low [66]. In the present study, the detection of Vitvi11g00497 gene, coding for auxin-responsive protein, in soft stage could suggest a possible role of this gene, as repressor of early auxin response genes at low auxin concentrations. Interestingly, this gene showed an expression below cut off in the study performed by [67], according to the Expression Atlas web tool (accessed June 8th 2021) and in the expression data obtained here (Figure S3). In a previous study in human research to identify potential causal genes with Crohn’ s disease, the risk alleles of these genes were found to be associated with lower expression levels of two target genes [68]. In this sense, eQTL works have frequently demonstrated that most eQTL effects are of relatively small size (<2-fold change in expression) [69, 17] which is in complete agreement with the results of large-scale experimental reports of putative regulatory variants [8].

Germin-like proteins (GLPs) are expressed in several developmental phases and organs in plants, as consequence of a number of abiotic and biotic stresses [70]. In grapes, the diversity present in the family of GLP has been studied, suggesting a role of VvGLP*3* in the defense response against one of the most devastating diseases of grapevines worldwide with a high economic impact, powdery mildew, produced by the Ascomycetous fungus Erysiph*e* necato*r* (syn. Uncinul*a necator*). In a fruit species such as plum (*Prunus salicina*), two novel GLP genes were isolated, exhibiting similar expression patterns throughout several phases of fruit development, except pit hardening and fruit ripening. According to the results, the accumulation of both Ps-GLP*s* is related to the evolution of auxin, since fruit development until the ripening stage, suggesting that auxin is affecting the regulation of both transcripts throughout the development and ripening of the fruits [71]. The identification of Vitvi14g02553 by Matrix eQTL in the early stage of fruit development (pre-véraison) in red cultivars, could suggest that this gene is putatively accumulated in the fruit during ripening in an auxindependent manner.

Another important identified gene family is Growth-regulating factors (GRFs). Originally, these plant-specific transcription factors, GRFS, were identified for their roles in leaf and stem development. More recently, some authors have highlighted them to be also important in key developmental stages including seed and flower formation, root development and the coordination of growth phases under adverse environmental conditions [72]. In *Arabidopsis thaliana*, GRFs are generally less expressed in mature ones than in actively growing tissues, with a function in initial phases of development and growth in different tissues [73, 74, 75]. The expression level of GRF Vitvi09g00107 was not high here, something that is correlated with the commented assumption about their high expression in early stages of growth.

In this study a gene coding for a PHD finger protein ING was Vitvi05g00925, having 8 genetic variants associated. This gene was only found in red cultivars in both stages and in the general analysis. This protein belongs to the ING family, having several proteins with conserved plant homeodomain (PHD)-type zinc fingers in their C-termini [76]. In this sense, a banana PHD-type, MaPHD1, which control their target gene expression by binding to the typical G-rich motif, having in the promoter a conserved core sequence of GNGGTG or GTGGNG [77, 78], seems to represses a cell wall-degradation gene MaXTH6 during fruit ripening. According to these authors MaPHD1 could directly suppress MaXTH6 which could be negatively associated, at least in part, with fruit ripening [79]. The results found in this study will be a solid foundation for a directed experiment to confirm the same negative association of Vitvi05g00925 with their target in grapes, which could bring more insights of grape ripening.

Other genes with a possible transcription activation activity are Vitvi06g00964, with 89 genetic variants associated by Matrix eQTL, and Vitvi06g00918, with 41 genetic variants associated. This last gene coding for a homeobox domain-containing protein, and was only observed in white cultivars in the soft stage by Matrix eQTL. Vitvi06g00964, with 89 genetic variants associated, was only found in red cultivars in both conditions and in both general analyses. This gene coding for a kinesin motor domain-containing protein. In this sense, novel functions have been attributed to kinesin motor proteins such as the regulation of gibberellin biosynthesis and cell elongation through transcriptional activation activity [80].

### 4.2 Conclusion

This study has provided new insights in grape ripening and manifests the advance of integrating different omics data for the study of complex process and traits. The accessibility to data underlying scientific publications allowed the author to efficiently apply this exhaustive methodology. Clearly, this study has exploited the value of mandated public data archiving (PDA) policies in the sciences, which is mandatory for top journals in several areas, including evolution and ecology. Overall, this work will improve the understanding of gene regulation during fruit ripening, since pre-véraison to post-véraison, in *Vitis vinifera* L. In the future, in addition of the detection of the lead eQTLs, the detected genes could be used for functional and mechanistic studies of grape ripening. Finally, and more importantly, the workflow developed here could be applied to other woody crops species, such as fruit trees or nut trees, where these types of studies are scarce and when the availability of such comprehensive data sets becomes a reality, which is not until to date.

